# *Osmia lignaria* (Hymenoptera: Megachilidae) produce larger and heavier blueberries than honey bees (Hymenoptera: Apidae)

**DOI:** 10.1101/2020.06.28.176396

**Authors:** Christine Cairns Fortuin, Kamal JK Gandhi

## Abstract

Fruit set, berry size, and berry weight were assessed for pollination by the solitary bee *Osmia lignaria* (Say) in caged rabbiteye blueberries (*Vaccinium ashei* Reade, Ericales : Ericaceae), and compared to that of uncaged rabbiteye blueberries which were pollinated largely by honey bees (*Apis mellifera* L). *O. linaria* produced berries that were 1.6mm larger in diameter and 0.45g heavier than uncaged blueberries. Fruit set was 40% higher in uncaged blueberries. This suggests that *Osmia* bees can produce larger and heavier berry fruit, but *O. lignaria* may be less efficient at blueberry pollination as compared to *A. mellifera* under field cage conditions.

---

Wild bees (Hymenoptera: Apidae) in the genera *Andrena, Bombus, Megachile, Osmia*, and *Peponapsis* make significant contributions to crop pollination (Winfree et al. 2008, Adamson et al. 2012). Wild bees are of particular value for crops such as alfalfa, blueberries, cranberries, oilseed rape, squash and tomato (Tepedino 1981, Cane 2002, Greenleaf and Kremen 2006, Ratti et al. 2008, Jauker et al. 2012), and significantly contribute to the diversity and sustainability of plants in natural systems (Fontaine et al. 2005, Tuell et al. 2014). Among wild bees, those in the genus *Osmia* are an economically important group of solitary cavity nesting bees that are often managed for tree pollination in various orchard systems including almond, apple, blueberry, cherry, and pear (Bosch and Kemp 2000, 2002, Monzón et al. 2004, Sampson et al. 2004, Bosch et al. 2006).

Wild bee species which commonly visit blueberries (*Vaccinium* spp.) include *Andrena* spp., *Bombus* spp., *Habropoda laboriosa*, and *Osmia* spp. (Sampson and Cane 2000, Tuell et al. 2009). *Osmia* spp. emerge concurrent with blueberry bloom in early spring and can be managed in orchard settings using nest blocks. Hence, *Osmia* bees can provide a viable alternative to *Apis mellifera* L. (honey bees), or compliment *A. mellifera* pollination in blueberry systems (Sampson and Cane 2000, Sampson et al. 2004). The presence of bumble bees (*Bombus* spp.) is correlated with larger berry size in blueberries and cranberries than those only associated with *A. mellifera* (Ratti et al. 2008), and the southeastern blueberry bee, *H. laboriosa*, has also been shown to improve blueberry size (Payne et al. 1989). Whether other wild bee species such as *Osmia* spp. can produce larger berries than *A. mellifera* has not been assessed in previous studies.

Rabbiteye blueberry (*Vaccinium ashei* Reade, Ericales : Ericaceae) is a species native to the southeastern United States (U.S.), and thus well adapted to the climate and growing conditions of this region. Rabbiteye blueberry tends to produce smaller fruits on average than other blueberry species, and therefore can be less competitive on the market against larger fruit producing species such as highbush blueberry (*Vaccinium corymbosum* L.) (Makus and Morris 1993). Rabbiteye blueberry is moderately to highly self-incompatible depending on the varietal, thus largely dependent on the presence of pollinators for adequate fruit set (Lyrene and Crocker 1983). One wild bee species which has been shown to be effective for management in rabbiteye blueberry is *Osmia ribifloris* (Cockerell), although *O. ribifloris* is not native to the southeastern U.S. where rabbiteye blueberry is grown (Sampson and Cane 2000). *Osmia lignaria* (Say) is a widely available commercial species which has been established as an effective managed pollinator for orchard crops such as apple, almond, and cherry (Torchio 1976, Kuhn and Ambrose 1984, Bosch and Kemp 1999, Bosch et al. 2000, Bosch et al. 2006), and the eastern subspecies (*O. l. lignaria*) is native to the temperate regions of the southeastern U.S. However, in highbush blueberry systems, *O. lignaria* has shown preference for other pollen resources, with pollen provisions exhibiting higher proportions of black cherry and clover than highbush blueberry (Pinilla-Gallego and Isaacs 2018). Thus, *O. lignaria* is generally considered to be a poor alterative for management in blueberry systems, however it is presently unclear if *O. lignaria* could be effective pollinators of rabbiteye blueberry if other “preferred” pollen resources are limited.

Our objectives were to determine the pollination efficiency of *O. lignaria* on rabbiteye blueberry, and determine if they are able to produce larger fruit than *A. mellifera*, as observed for other wild bee species such as *Bombus* spp (Ratti et al. 2008). Fruit set, berry size, and berry weight were assessed for *O. lignaria* in caged rabbiteye blueberries, and compared to those of uncaged blueberries which were pollinated largely by *A. mellifera*.

## Materials and Methods

*O. l. lignaria* adult cocoons were obtained from a commercial source (Mason Bee Company, Deweyville, Utah) in February 2019 and were kept in cold storage at 4^°^C until ready for emergence. Before emergence, cocoons were washed in 5% (by volume) bleach solution, and rinsed with clean water to remove any contaminates or mold. Four hundred cocoons were emerged in the laboratory at ambient temperature (25°C) between 11-14 March 2019. Emerged bees were returned to cold storage until ready for use. Once 120 females and 180 males had emerged successfully, the adults were randomly distributed into groups of 20 females and 30 males per group.

Six (6 × 6 x 2 m^2^) flight cages were set up in the permanent blueberry orchard at Durham Horticulture Research Farm, University of Georgia in Watkinsville, Georgia. The orchard contains alternating rows of Climax and Premier varietals of blueberries. Each flight cage enclosed 6-7 mature Rabbiteye blueberry shrubs, with a minimum of three shrubs of each varietal within each cage. Cages were erected during stage 4 of bud development (Spiers 1978), before the opening of flowers. Groups of 20 female and 30 male *O. lignaria* were released on 21 March 2019 into each cage. Each cage contained a Styrofoam nest block with 36 nesting holes filled with 8mm diameter paper tubes, and a 15 × 15 cm^2^ mud pit was created and maintained within each tent as nest material for the female bees.

Five blueberry shrubs inside each cage and five blueberry shrubs outside each cage were chosen for the study. Racemes on shrubs were marked during stage 5 of bud development with metal tags after cages were set up. Three racemes on each of the five shrubs inside the cages, and three racemes on each of the five shrubs outside the cages were selected and marked in the same way, for a total of 15 marked racemes for each cage or uncaged area. The total number of florets on each terminal bud were marked on the tag. Cages were set up before the opening of the blueberry flowers, and removed well after fruit drop, and therefore, berries inside the cages were exclusively pollinated by *O. lignaria*. A honey bee apiary consisting of six hives was situated about 50 m from the blueberry patch, and thus pollination of blueberries outside of the cages was largely accomplished by *A. mellifera* (though wild bees could have played some role in pollination of uncaged plants).

Fruit set was counted on 24 April 2019 while berries were in their late green fruit stage, comparing labeled number of florets to set fruit. On 10 June 2019, 20 berries were harvested at random from each marked shrub. The harvested blueberries were then weighed in their groups of 20, and a subset of 10 berries from each group was measured to the nearest 0.1mm using a caliper. Berries were measured along the widest circumference. Randomness of berry selection for measurement was ensured by using a random number generator after arranging berries in a line using 1-20 numbers. *Osmia* bees had to be pulled from the cages after 21 days due to unforeseen circumstances, thus the blueberry shrubs inside of the cages did not receive a full bloom cycle of pollination, which continued for two more weeks after removal.

All analyses were conducted using R statistical software (RStudio Team 2019). Berry weight and berry size were compared between the caged (*O. lignaria*-pollinated) and uncaged (*A. mellifera*-pollinated) blueberries using a two-way ANOVA with each caged and adjacent uncaged area treated as a block for a total of six blocks. Assumptions of normality and constant variance were checked by examination of normal Q-Q and residuals vs. fitted value plots. Each set of 20 harvested berries was used as the replicate for weight and size (six mean measurements for each cage and for each uncaged area, n=36 caged, n=36 uncaged). For weight, the combined weight of each set of 20 berries was divided by 20 to obtain the average weight per blueberry for that set. For size, the average size measurement of ten randomly selected blueberries from each set of 20 was used as the replicate measurement.

Differences in fruit set between caged and uncaged flowers were analyzed using a generalized linear model with negative binomial distribution utilizing the MASS package in R (Venables and Ripley 2002). Some metal tags marking racemes were lost in a windstorm, particularly effecting the uncaged tags. Thus, ultimately there were n=104 fruit set measurements in cages and n=62 measurements in uncaged areas. Each raceme’s measurement was treated as the unit of replication, and each caged and adjacent uncaged area treated as a block, for a total of six blocks. Fishers’ F-test was used to determine equality of variances. The response variable for the model was the total number of set (ripe) fruit on each marked raceme, offset by the number of original florets on the raceme times the number of weeks pollinators were present, which was three weeks for inside the cages, five weeks outside of the cages.

## Results

Caged berries pollinated by *O. lignaria* were on average 1.6 mm larger in diameter and 0.45g heavier than uncaged berries pollinated largely by *A. mellifera* (weight: *F*_1,65_ = 56.3, *P* < 0.001; size: *F*_1,65_ = 92.3, *P* < 0.001), a difference which was clearly visible to the naked eye (Figs. 1-2). Uncaged blueberries averaged (±SE) 14 ± 0.03 mm in size and weighed 1.44 ± 0.08 g. Blueberries pollinated by *O. lignaria* averaged 15.6 ± 0.02 mm in size and weighed 1.85 ± 0.09 g. Berry weight varied between blocks (weight: F_5,65_ = 2.6, p = 0.035; size: F_5,65_ = 2.2, p = 0.069), but *O. lignaria* pollinated blueberries were larger than their uncaged counterparts within all blocks (Fig. 1). Fruit set of uncaged shrubs was on average 40% higher than for caged and this difference was significant (*Z* = 1.98, *P* = 0.048) [Figure 3].

**Fig. 1.**
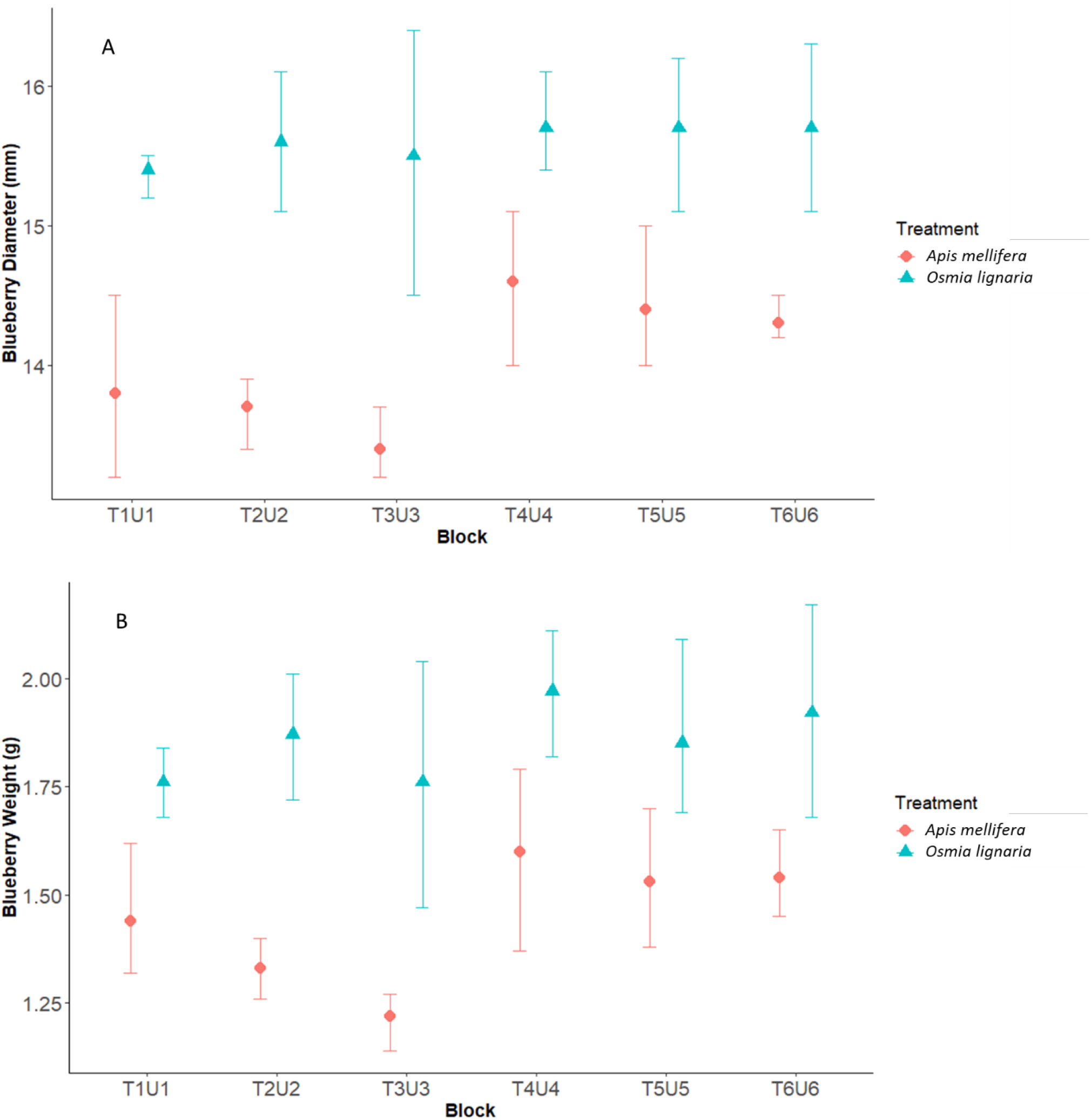
Means and 95% confidence intervals of blueberry diameter (A) and weight (B) measurements of blueberries pollinated by *Osmia lignaria* (triangles) vs. those pollinated primarily by *Apis mellifera* (circles) by blocks (i.e., T1U1 = Tent 1 and the Uncaged area adjacent to Tent 1, etc.).

**Fig. 2.**
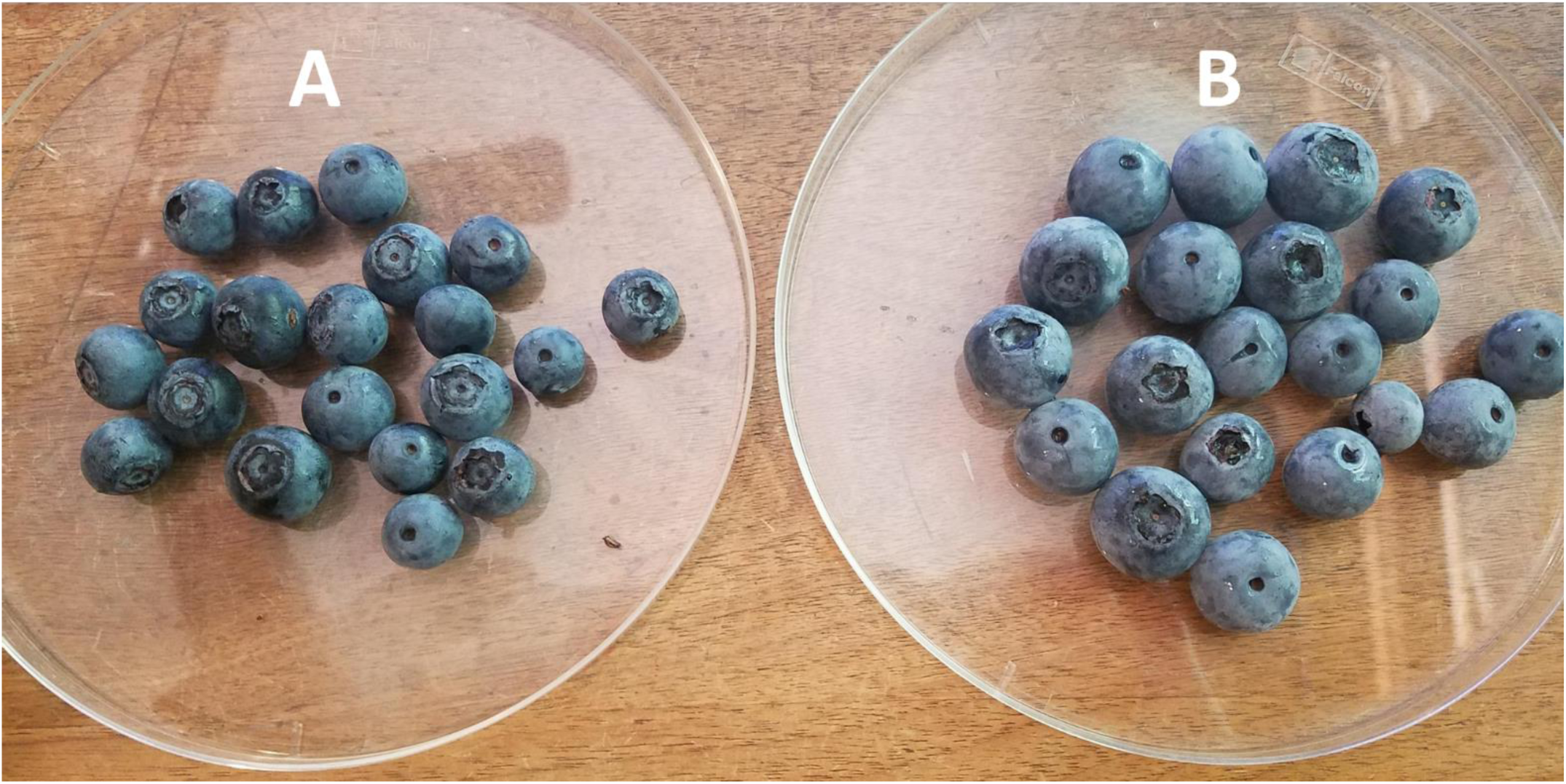
Example of one set of 20 blueberries harvested from shrubs pollinated by *Apis mellifera* (A) and *Osmia lignaria* (B).

**Fig. 3.**
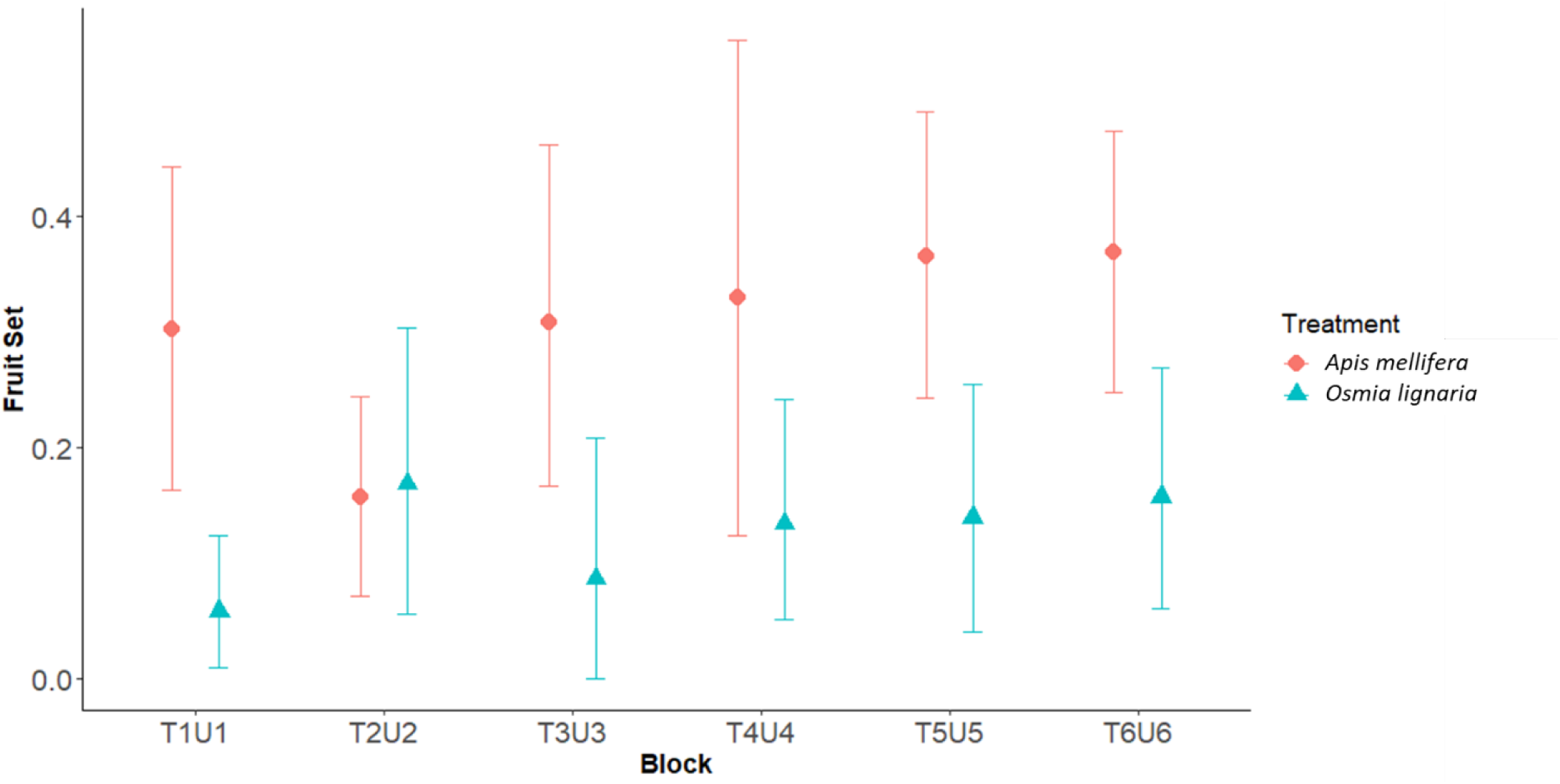
Fruit set (percentage of florets on terminal bud setting fruit) mean and confidence intervals for the caged *Osmia lignaria* (triangles) compared to uncaged pollination by *Apis mellifera* (circles) by blocks (i.e., T1U1 = Tent 1 and the Uncaged area adjacent to Tent 1, etc.).

## Discussion

The current study results suggest that *Osmia* spp. have the potential to produce ∼11% larger and ∼22% heavier blueberry crops, although *O. lignaria* may be less efficient as compared to *A. mellifera* for blueberry pollination. Increased opportunity for outcrossing of pollen in open areas primarily by *A. mellifera* may be a contributing factor to the 40% higher fruit set for uncaged plants. Previous studies have found fruit set of rabbiteye blueberry pollinated *O. ribifloris* to be comparable to that of *A. mellifera* (Sampson and Cane 2000), and estimated economic returns from a single female *O. ribifloris* to be between $12 – $24 (Sampson et al. 2004). If *O. lignaria* is considered a surrogate for *O. ribifloris* and other bees, our current study suggests that in addition to effective pollination, *Osmia* bees may also produce larger berries.

There could be effects of cages on pollination efficiency, however other studies such as on *Osmia cornifrons* (hornfaced bees) have reported when various species of bees were individually caged with blueberries, they contributed to pollination at the same level as *A. mellifera* (West and McCutcheon 2009). Future studies may consider comparisons of fruit size for flowers pollinated by *Osmia* spp. and *Apis* spp. in natural settings. It is possible that *O. lignaria* may disperse to find more preferred resources when they are not caged. However, for *Osmia* bees such as *O. ribifloris* which have been shown to prefer blueberry, we suggest that these bees may be highly beneficial for rabbiteye blueberry size and quality, and hence their management and inclusion in agricultural systems merits further investigation (Bosch and Kemp 2002).

## Acknowledgements

This work would not have been possible without contributions of the following individuals and laboratories: Keith Delaplane and staff of the University of Georgia (UGA) College of Agriculture and Environmental Sciences Honey bee Lab; Ryan McNiel and staff the UGA Durham Horticulture Research Farm (UGA, College of Agriculture and Environmental Sciences); Berry Brosi (Emory University); Elizabeth McCarty (UGA, D.B. Warnell School of Forestry and Natural Resources); Diana Cox-Foster, Ellen Klomps, and Theresa Pitts-Singer (USDA ARS Pollinating Insects Lab); Natalie Boyle (Penn State University, College of Agricultural Sciences); Brittany Barnes, Lea Clark, and Michael Del Rossi (UGA, D.B. Warnell School of Forestry and Natural Resources, Forest Entomology Lab), and John Fortuin and Pumpkin. Funding was provided by the USDA Southern SARE grant, EPA STAR Fellowship, and UGA, D.B. Warnell School of Forestry and Natural Resources.

